# A predictive unifying explanation for nuclear shapes based on a simple geometric principle

**DOI:** 10.1101/2022.09.30.510302

**Authors:** Richard B. Dickinson, Tanmay P. Lele

## Abstract

Nuclei have characteristic shapes dependent on cell type, which are critical for proper cell function, and nuclei lose their distinct shapes in multiple diseases including cancer, laminopathies, and progeria. Nuclear shapes result from deformations of the sub-nuclear components—nuclear lamina and chromatin. How these structures respond to cytoskeletal forces to form the nuclear shape remains unresolved. Although the mechanisms regulating nuclear shape in human tissues are not fully understood, it is known that different nuclear shapes arise from cumulative nuclear deformations post-mitosis, ranging from the rounded morphologies that develop immediately after mitosis to the various nuclear shapes that roughly correspond to cell shape (*e.g*., elongated nuclei in elongated cells, flat nuclei in flat cells). Here we establish a simple geometric principle of nuclear shaping: the excess surface area of the nucleus (relative to that of a sphere of the same volume) permits a wide range highly deformed nuclear shapes under the constraints of constant surface area and constant volume, and, when the lamina is smooth (tensed), the nuclear shape can be predicted entirely from these geometric constraints alone for a given cell shape. This principle explains why flattened nuclear shapes in fully spread cells are insensitive to the magnitude of the cytoskeletal forces. We demonstrate this principle by predicting limiting nuclear shapes (i.e. with smooth lamina) in various cell geometries, including isolated on a flat surface, on patterned rectangles and lines, within a monolayer, isolated in a well, or when the nucleus is impinging against a slender obstacle. We also show that the lamina surface tension and nuclear pressure can be estimated from the predicted cell and nuclear shapes when the cell cortical tension is known, and the predictions are consistent with measured forces. These results show that excess lamina surface area is the key determinant of nuclear shapes, and that nuclear shapes can be determined purely by the geometric constraints of constant (but excess) nuclear surface area and nuclear volume, not by the magnitude of the cytoskeletal forces involved.

## 1. Introduction

The shape of the mammalian cell nucleus is an important cellular feature that varies in different cell types and tissues. For example, in endothelial cells lining the blood vessels and capillaries, nuclei are typically flat, whereas they are more rounded in epithelia and elongated in fibroblasts. Although the mechanisms regulating nuclear shape in human tissues are not fully understood, it is known that different nuclear shapes arise from cumulative nuclear deformations post-mitosis, ranging from the rounded morphologies that develop immediately after mitosis to the various nuclear shapes that roughly correspond to cell shape (*e.g*., elongated cells have elongated nuclei, flat cells have flat nuclei). The cell nucleus is deformed by mechanical stresses generated in the cytoskeleton[1], and the resulting nuclear deformations are critical for mediating essential cellular activities. In particular, nuclear deformations enable fibroblast migration during wound healing[2], cell migration during early embryonic differentiation[3], neuronal migration[4] and neurokinesis during development[5], immune cell migration across the endothelium[6], and muscle contraction[7].

In addition, nuclear deformations can trigger cell-type-specific gene expression and signaling pathways through mechanisms that are currently not well-understood, due to an inadequate understanding of how sub-nuclear structures respond to nuclear forces. However, it has been shown that stretching of the nuclear envelope induces the opening of stretch-activated ion channels in the nuclear membrane, which in turn, activates the small GTPase RhoA and alters cell migration activity[8]. Similarly, mechanical flattening of the nucleus can stretch nuclear pores, leading to translocation of yes-associated protein (YAP) into the nucleus[9; 10], and thereby alter gene expression[11].

Critically, nuclear deformations can also have pathological consequences. For example, extreme deformations may tear and rupture the nuclear envelope, leading to DNA damage, tumorigenesis[12], and invasive migratory phenotypes[13]. Additionally, nuclear deformation due to a decrease in nuclear lamin A levels allows cell migration through confining spaces in the tissue interstitium, thereby contributing to tumor cell escape and metastasis[14; 15; 16]. Cancer pathologists commonly use nuclear morphology to grade different cancers, assessing both shape deformation and enlargement in size (reviewed by us in [17]).

Generally, nuclear shape mimics the overall cell shape. For example, nuclei are elongated in elongated cells, and they take on a flattened disk-like shape in well-spread cells [18; 19; 20]. The predominant model for explaining such shapes assumes that the nuclear lamina, chromatin, and other sub-nuclear structures deform elastically in response to cytoskeletal forces (reviewed previously in [11]). In this model, the nucleus is assumed to be a stiff, elastic object that deforms from an initial spherical shape by elastically straining the chromatin and stretching the stiffest surface element, which is the nuclear lamina. The models are motivated by measurements of nuclear deformation with force probes, such as the Atomic Force Microscope (AFM) and other techniques (*e.g*., micropipette aspiration) that apply controlled forces on short timescales of only a few seconds[21; 22; 23; 24; 25; 26; 27; 28; 29; 30; 31; 32; 33]. In these models, the resting undeformed state of the nucleus is commonly assumed to be a sphere with a thin, mechanically stiff, surrounding lamina layer. However, since a sphere is the unique geometric shape with a minimal surface area for its volume (or maximal volume for its surface area), any deformation of a sphere requires either a change in surface area, a change in volume, or both. Consequently, if the resting nucleus is assumed spherical, the large deformations such as observed in flattened or elongated nuclei, would require large enough forces on the nucleus to either stretch the stiff nuclear lamina or to compress the nuclear volume. Yet, three-dimensional reconstructions of the nuclear shape in rounded cells show that the lamina is not a smooth spherical shell; rather, it has a significantly greater surface area than that of a sphere of the same nuclear volume, with the excess lamina area stored in surface folds, wrinkles, and undulations [20; 34; 35; 36; 37] (Figure 1A). When an object’s surface area exceeds that of a sphere of the same volume, a wide range of three-dimensional shapes are geometrically possible for that volume and (excess) surface area without requiring a mechanical stretching of the surface. We have previously invoked excess lamina surface area to explain why the nucleus is highly compliant to shape changes during spreading of cells, and only is “stiff” to further flattening once the surface area is smoothed and where further deformations require volume compression or lamina area expansion[20]. Our observed correlation between cell spreading and nuclear flattening, as well as the asymptotic limit to nuclear flattening, did not depend on various perturbations of the cytoskeleton including myosin inhibition. Further, our prior experiments with migrating fibroblasts show that removal of cytoskeletal stresses does not result in relaxation of elongated nuclear shapes back to circular (spherical) shapes[38]. Thus, contrary to the dominant mechanical models for nuclear shaping in the literature, we concluded that the asymptotic, limiting nuclear shape in fully spread cells does not reflect a balance between elastic stresses in the nuclear shape and cytoskeletal forces, and that any elastic energy that would cause the nucleus to recover its initial shape dissipates on the time scale (minutes) of cellular and nuclear shape changes.

**Figure 1.**
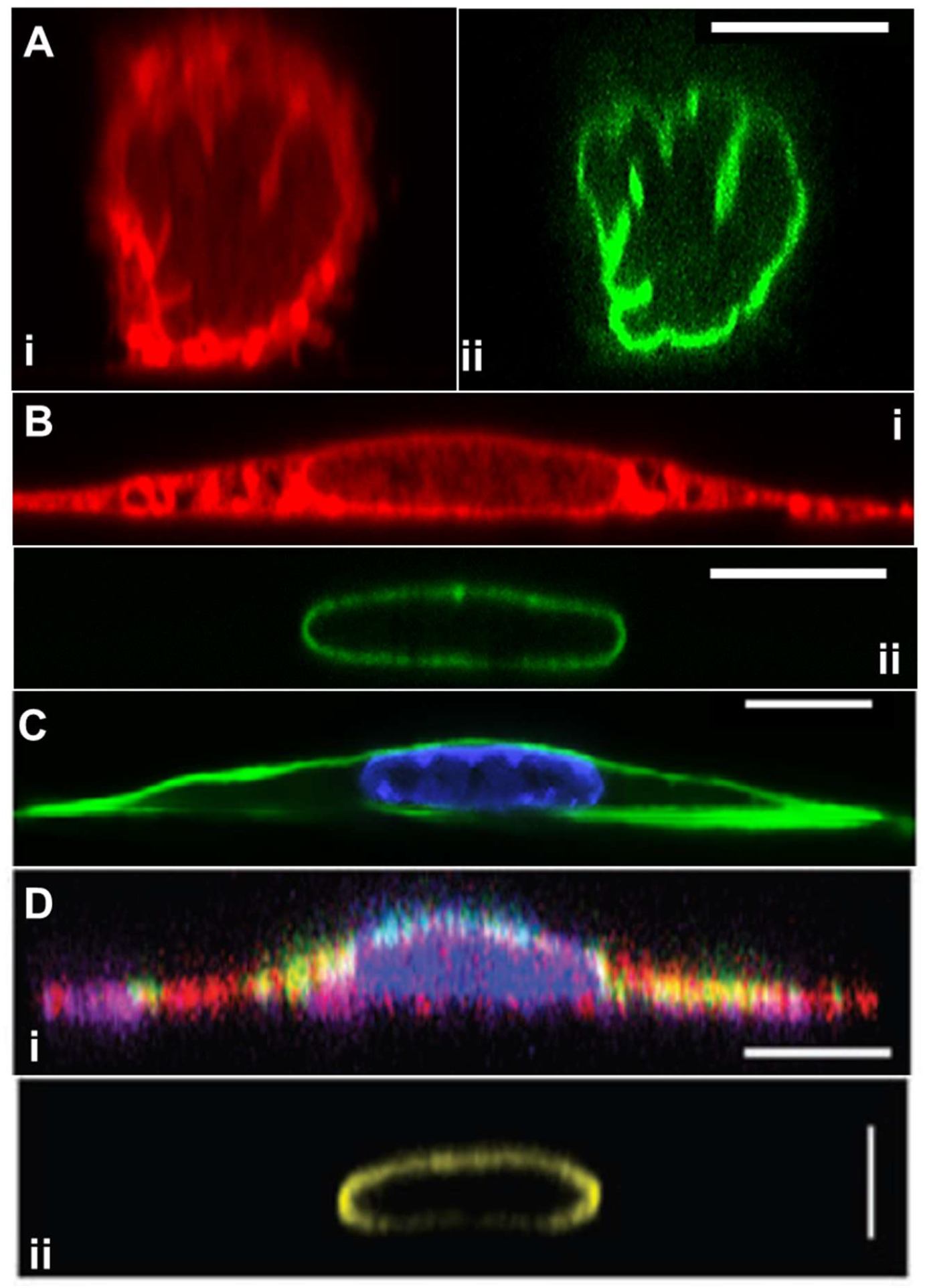
Geometry of rounded and fully spread cells from x-z perspective. (A.) NIH3T3 fibroblasts fixed within five minutes of surface attachment with two fluorescence channels: (i) labeled with Dil D7556 lipid dye or (ii) GPF lamin-A (image from Dickinson et al. *APL Bioeng* 2021 [34]). Excess area is seen clearly in the form of surface folds, wrinkles, and undulations. (B). Same conditions as (A) but at 24 hrs of spreading, showing different apical and side curvatures of smooth nuclear lamina, and the different apical cell surface curvatures on the nuclear cap versus elsewhere. (C) Another x-z perspective of a NIH3T3 fibroblast where f-actin is labeled with phalloidin (green) and the chromatin is labeled with Hoechst (H33342), clearly showing the cortical actin (from Katiyar et al. *Soft Matter* 2019 [35]). (D) An x-z perspective of a myoblast cell and lamina from Jana et al. *Adv Sci* 2022 [39] with (i) labeled cytoskeleton (actin (red), tubulin (purple), and vimentin (green)) and (ii) labeled lamin A/C. All scale bars are 10 microns.

Motivated by the above observations, we show in this paper that the limiting nuclear shapes with smooth (tensed) lamina can be closely predicted based solely on a simple geometric principle: the excess surface area of the nucleus (relative to that of a sphere of the same volume) permits a wide range highly deformed nuclear shapes under the constraints of constant surface area and constant volume, but when the lamina is smooth (tensed), the nuclear shape can be predicted entirely from these geometric constraints alone for a given cell shape. By accounting for the excess lamina surface area, highly deformed nuclear shapes can be predicted in various geometries without invoking further mechanical principles such as elastic deformation in response to cytoskeletal forces. Indeed, the observed nuclear shapes can be geometrically predicted independent of the magnitude of the cytoskeletal forces involved.

## 2. Theoretical Model

The unique shape with a minimum surface area under the constraint of constant volume is a sphere. Its surface area *A*_sphere_ is geometrically related to its volume *V* by

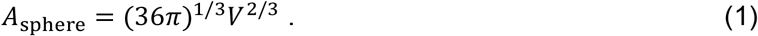

Because its area is at minimum, any deformation of shape from the sphere requires an increase in surface area, a decrease in volume, or both. A well-known physical example of a sphere in nature is a drop of water in oil which takes on a spherical shape to minimize its surface energy, equal to the product of surface tension and surface area. However, unlike a liquid drop, the nuclear lamina in the surface of a rounded nucleus is not spherical; rather it exhibits surface folds, wrinkles, and undulations, which only disappear in fully spread cells (Figure 1 A,B). This implies that in the unstressed nucleus, the lamina surface area, *A*, is greater than *A*_sphere_, with a fractional excess surface area *ε* defined as

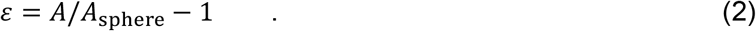

Unlike the unique shape of a sphere with volume *V* and surface area *A*_sphere_, an object with an excess surface area *A* > *A*_sphere_ may take on a wide range of possible shapes with the same volume and surface area. Consequently, specifying only the constraints of fixed volume and fixed (but excess) surface area is insufficient to determine a unique shape. To illustrate this, we examine an object’s shape with fixed volume and fixed surface area confined in a cylindrical pore (Figure 2A). For a given pore radius, there is a unique shape, a capsule, with cylindrical sides and hemispherical endcaps that has a minimum surface area (at *A*_min_) for a given volume, *V*. Therefore, a shape with fixed volume *V* and fixed surface area *A*, can only fit within the pore if *A* ≥ *A*_min_; otherwise, if *A* < *A*_min_, the pore is too small to accommodate the fixed surface area. When the pore diameter is at the threshold size where *A* = *A*_min_, then the object must take on the unique limiting shape of a capsule. Figure 2A shows capsules with a fixed volume and varied values of the fractional excess area *ε* at the threshold pore diameter. When *A* > *A*_min_, the shape within the pore is not completely determined by the volume and area constraints, since it could take on any number of shapes. For example, it could be even narrower, have non-spherical endcaps, or additional surface area that could be shaped into various surface folds, wrinkles undulations, etc. Since migrating cells in the body may find themselves in such cylindrical confinements in different contexts [12; 15; 40], the biological implication is that a cell nucleus with incompressible volume and excess surface area has no geometric constraints to moving through a pore above the threshold diameter, but it would have to either stretch its lamina area or compress its volume to move through a smaller pore.

**Figure 2.**
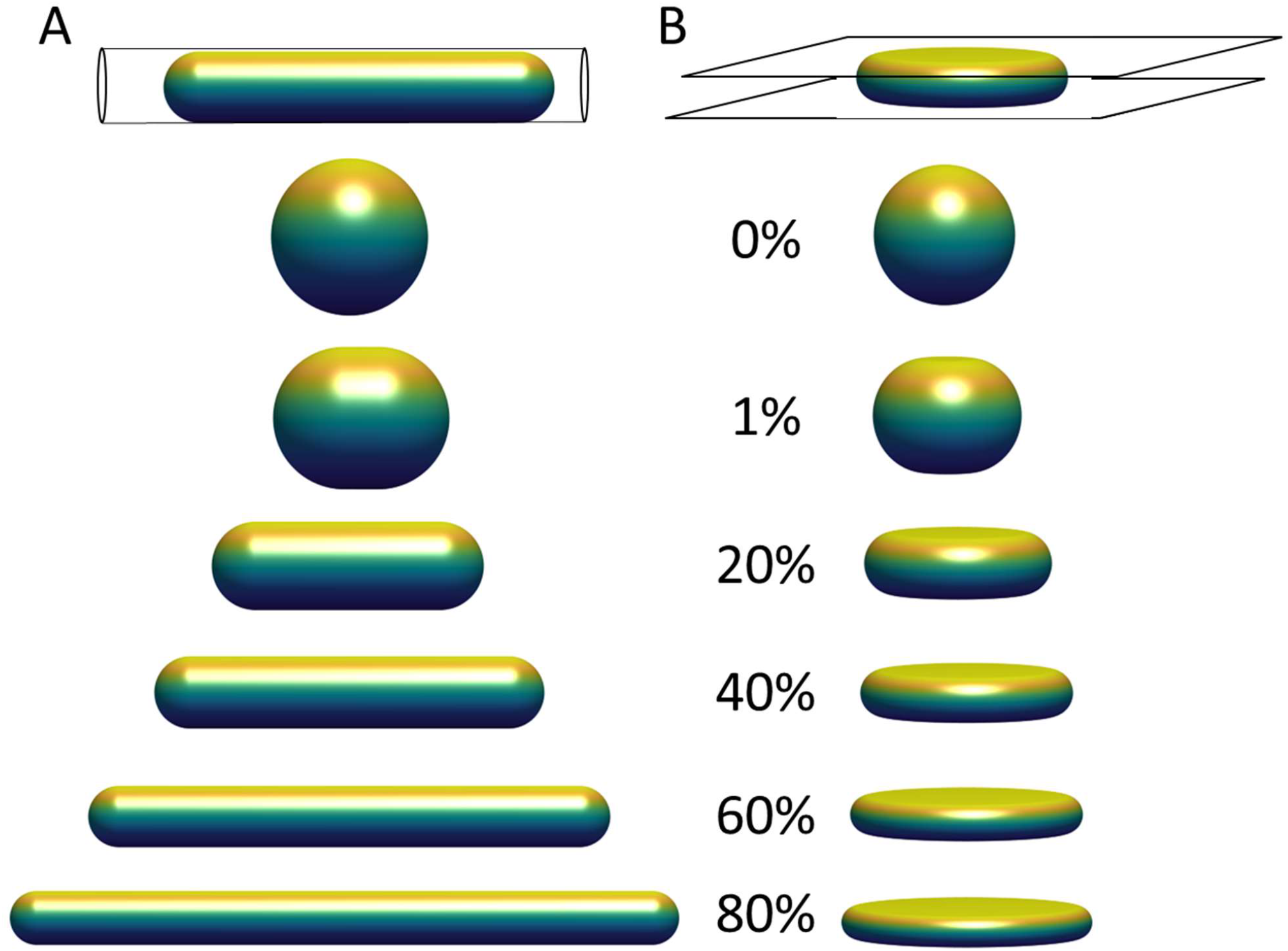
Limiting shapes in confined geometries of constant volume and constant surface area, with increasing fractional excess surface area relative to that a sphere of the same volume. Limiting shapes are shown for objects confined to (A) a cylindrical pore; and (B) between two flat plates. The excess surface area allows the object to occupy a smaller length dimension of confinement (pore diameter or gap distance between the plates), and the shapes shown represent the unique shape at the threshold length, below which there are no shapes possible that satisfy the fixed volume and area constraints, and above which infinite shapes are possible (e.g., with surface folds, wrinkles, or undulations).

Another relevant example is an object with constant surface area and constant volume vertically compressed between two flat surfaces (Figure 2B). Similar to Figure 2A, there is a threshold gap distance for an object confined between the two plates above which infinite shapes are possible for a fixed volume and excess area, but no shapes that satisfy these constraints are possible at smaller gap distances. At the threshold gap distance, the unique shape that satisfies the geometric volume and area constraints is a disk-like shape with nodoid sides. Similar to the hemispherical surfaces of a capsule, nodoids are convex axisymmetric surfaces with constant mean curvature[41], and they have a minimum surface area for the enclosed volume. If the object only resists changes to volume and surface area, maintaining a flattened shape would require no force at gap distances above the threshold gap distance. This would explain why nuclei in spreading cells asymptotically reach the same minimum height even when myosin activity is inhibited [20], i.e. flattening the nucleus down to the minimum height can proceed without changing nuclear volume or lamina area while requiring minimal force, but flattening the nucleus below the minimum would stretch the (now taut) stiff lamina or compress the nuclear volume, which requires much higher force and is not observed during spreading. In these examples, the excess surface area permits a wide range of possible shapes without necessitating any compression of the volume or expansion of the surface area, but only down to a certain threshold length (i.e., pore size or gap distance) of confinement. For example, the cell nucleus can store its excess area in various surface folds, wrinkles, and undulations, which become smoothed out only when a flattened height is reached during cell spreading. This transition is clearly observed when the wrinkled lamina in rounded cells becomes smoothed out when the nucleus flattens in fully spread cells (Figure 1; [34]).

In both examples in Figure 2, the surfaces that are not in contact with the walls are surfaces of constant curvature (hemispherical caps in Figure 2A or nodoid sides in Figure 2B). When there is a surface tension, then the stress balance requires a corresponding pressure difference Δ*P* across the interface. The surface tension *τ* of an interface is related to Δ*P* and the mean surface curvature *H* by Laplace’s Law, i.e.

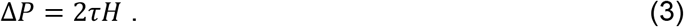

Though *H* is known from the shapes, Eq. 3 indicates that *τ* and Δ*P* cannot be obtained independently from the curvatures alone (only their ratio can be obtained), reflecting the fact that the limiting shapes with surfaces of constant mean curvature result from geometric constraints rather than the magnitude of the forces. That is, the same limiting shape is generated regardless of overall magnitude of the forces involved. This principle is consistent with the observation that the limiting heights of flattened nuclei in spread cells do not depend on the presence of specific cytoskeletal structures or myosin activity[20], provided the cell is able spread enough to vertically confine the nucleus to its limiting shape. Hence, the geometric constraints permit the same highly deformed nuclear shapes regardless of the magnitude of the force, and when the lamina is smooth, the pressure and surface tension are related to each other by the surface curvature.

## 3. Application of Model to Interpret Observed Nuclear Shapes

### 3.1 Nuclear shapes in a cell spreading on a flat substratum

We now extend the above conclusions to nuclear shapes in spread cells confined to various geometries to show that deformed, limiting shapes can be predicted for a given (excess) surface area, nuclear volume, and cell volume. This calculation attempts to simultaneously capture the asymptotically limiting cell and nuclear shapes in spread cells, and it neglects additional stresses that may arise from movement of the cell boundary or other cellular shape changes such as cell crawling [20; 35; 38]. That is, we draw a distinction between the viscous or viscoelastic forces that drive the nucleus toward the limiting shape and any forces that may be required to sustain the limiting shape, focusing on the latter. First, we consider the case of an axisymmetric cell spread on a flat surface, where the nuclear and cell shape can be solved analytically (see derivation in Methods, Section 5). Cell and nuclear shapes were calculated by solving for the surfaces of constant mean curvature that satisfy the constraints of fixed lamina area, cell volume, and nuclear volume. The relevant interfaces are the cell cortex interface with the surrounding medium (an unduloid surface of curvature *H*_*cell*_), the nucleus-cytoplasm interface (a nodoid surface of curvature *H*_*nuc*_), and the joint nucleus-cortex interface with the surrounding medium (a spherical cap of curvature *H*_*cap*_). These three distinct regions of different curvatures are commonly seen in x-z cross-sections of fully spread cells (see, for example, Figure 1B-D). Similar to the shape calculation in Figure 2B, increasing values of excess area generates limiting nuclear shapes that are increasingly flattened against the substratum (Figure 3). Moreover, the calculated shapes are quite similar to experimentally observed x-z profiles of cell and nuclear shapes in fully spread cells (compare with Figure 1), which regularly show three curved surfaces for the nuclear cap, cortex, and nuclear sides that are similar to the corresponding calculated surfaces. This agreement based on geometric considerations alone supports our assertion that the flattened limiting shapes of nuclei observed in spread cells are geometrically determined, independent of the magnitude of the cytoskeletal forces involved.

**Figure 3.**
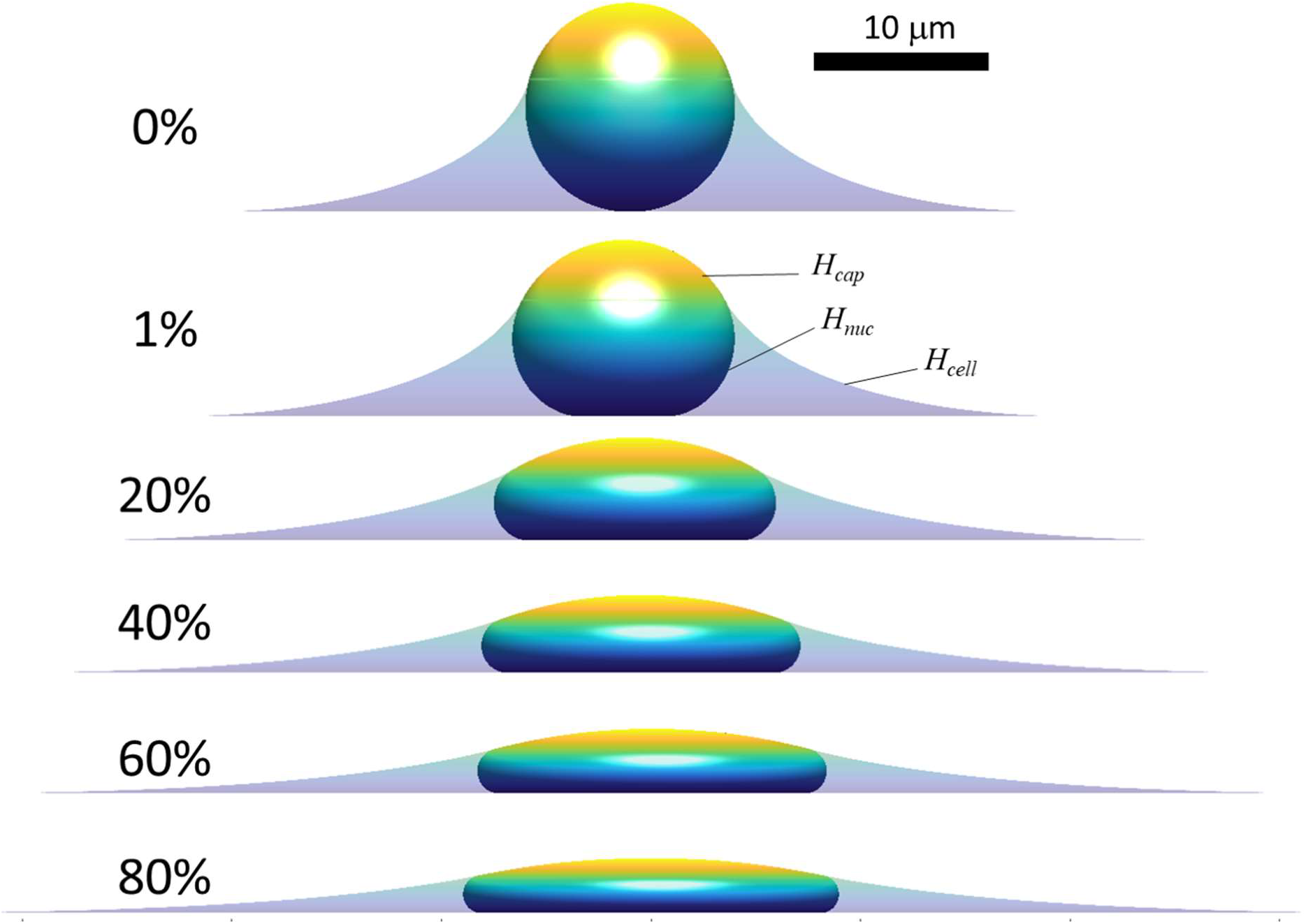
Calculated axisymmetric cell and nuclear *x*-*z* profiles in spread cells are shown for increasing amounts of fractional excess areas indicated by the percentages. Like experimental x-z profiles such as those shown in Figure 1, spread cells with smooth nuclear lamina have surfaces of constant mean curvature due to pressure differences across the three different interfaces, including the cortical cell surface (mean curvature, *H*_cell_), the nuclear surface (containing the lamina) in contact with the cytoplasm (*H*_nuc_), and the joint cortex/nuclear apical surface capping the cell (*H*_cap_.) The cell and nuclear shapes were found by solving for the mean curvatures of each surface that satisfy the constant volume and surface area constraints (see Methods). For the profiles shown here, the spreading cell radius was taken as the radius where the slope of the cell edge just becomes parallel to the substratum. As shown, increasing excess area permits flatter nuclei and greater spreading radius for the same nuclear and cell volume (*V* and *V*_cell_, respectively). For the calculations shown here *V =* 900 µm^3^, *V*_cell_ = 3.4*V*, consistent with values for NIH 3T3 fibroblasts[20]. Comparison to experimental profiles like in Figure 1 suggests nuclei typically have 30-60% excess surface area, which allows a flat equilibrium nuclear shape that is geometrically determined and independent of the magnitude of cellular forces.

When the lamina is smoothed and tensed, the surface curvatures are related to the surface tensions and pressures by Laplace’s Law. Following Eq. 3, the stress balances across the cortex-cytoplasm, cortex-nucleus and nucleus-cytoplasm interfaces are:

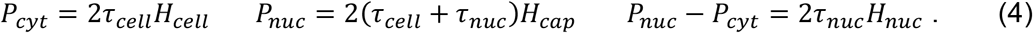

where the pressures defined are relative to the surrounding pressure. Rearranging Equation 4 yields

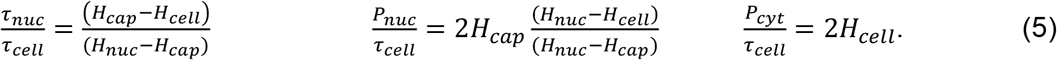

Hence, *P*_*nuc*_, *P*_*cyt*_, and *τ*_*nuc*_ can be calculated from the measured or calculated surface curvatures when *τ*_*cell*_ is known. Eq. 5 implies again that the shapes alone, which are geometrically constrained limits, do not the yield the overall force magnitude, only the relative tensions and pressures. But, when the cortical tension *τ*_*cell*_ is known, the nuclear and cytoplasmic pressures and the lamina tension can be calculated. Figure 4 shows calculations for the curvatures, pressures, and lamina surface tension for increasing values of excess lamina surface area at the same cell spreading radius and a specified value of *τ*_*cell*_ = 0.5 nN/µm [42]. As shown in the table, increasing the excess lamina area is predicted to correspond to a decrease in nuclear pressure and surface tension for the same spreading radius. For a larger values of excess area, the solution to the cell and nuclear shapes has a negative lamina tension, which is assumed non-physical. In this regime, rather than conforming to the apical surface, the nucleus would instead be expected to separate from the cortex and take on any of the range of possible shapes that would satisfy the constant volume and surface area constraints within the gap between the spherical-cap shaped cortical surface and the substratum. Generally, the geometric constraints of constant area, nuclear volume, and cell volume only allow unique solutions with a positive lamina tension (*τ*_*nuc*_ > 0, *P*_*nuc*_ > *P*_*cyt*_) above a certain cell spreading radius, where the nuclear pressure and lamina tension can be calculated. Below this radius, though, infinite solutions with *τ*_*nuc*_ = 0 and *P*_*nuc*_ = *P*_*cyt*_ are possible (e.g., with lamina folds and wrinkles). This range is illustrated further in Figure 5, where the cell shapes, surface curvatures, and pressures are calculated for the case of 40% excess lamina area, over the range of cell spreading radii that permit solutions with non-negative lamina surface tension (*τ*_*nuc*_ ≥ 0). At the smallest radius, where *τ*_*nuc*_ = 0, the nucleus conforms to the spherical-cap-shaped cell and the nuclear and cytoplasmic pressures are equal. At maximum spreading, taken to be where the cell edge becomes parallel to the substratum, the lamina surface tension is largest.

**Figure 4.**
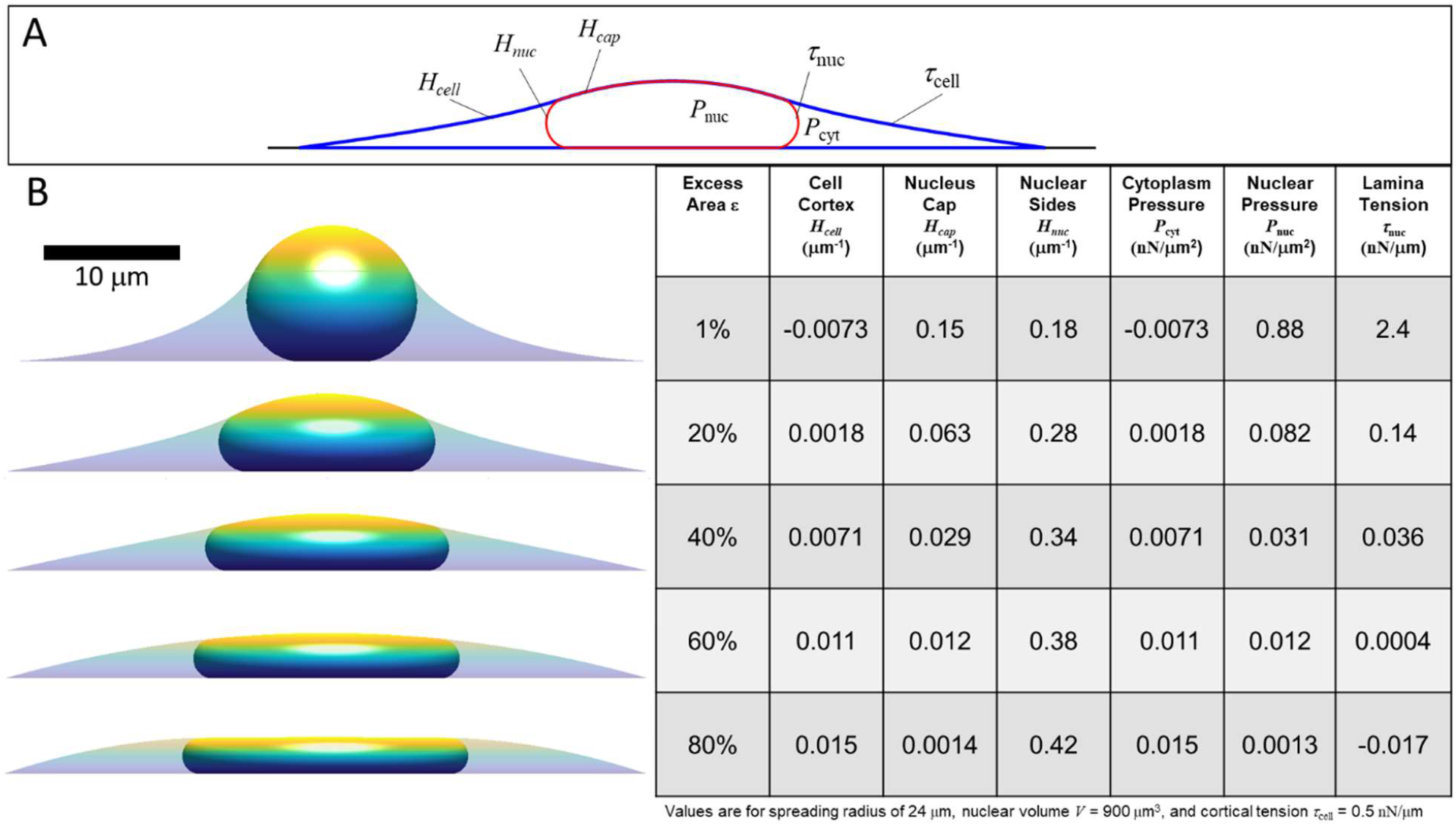
Calculated axisymmetric nuclear shapes showing the predicted effect of excess lamina area on the values of lamina surface tension, nuclear pressure, and cytoplasmic pressure, for a given cell spread radius. (A) The three curved interfaces are (1) the cell cortex (mean curvature *H*_cell_ and surface tension *τ*_cell_), (2) nuclear interface with the cytoplasm (mean curvature *H*_nuc_ and tension *τ*_nuc_), and (3) the apical cap where the lamina and cortex are in contact, interfacing the nucleus and the surrounding media (curvature *H*_cap_ and net surface tension equal to *τ*_nuc_ + *τ*_cell_) (B) Cell and nuclear shapes are shown for the same spread radius and increasing amounts of excess surface area shown in the adjacent table column. The table shows the calculated curvatures, pressures, and lamina tension based on the excess area and the nuclear and cell volume constraints (in this case, *V =* 900 µm^3^, *V*_cell_ = 3.4*V*, consistent with values for NIH 3T3 fibroblasts[20]. The force scale is set by assuming *τ*_cell_ = 0.5 nN/µm. Note that for 80% excess area at this cell spreading radius, the calculation yields a negative lamina tension, which is a non-physical solution implying the spread radius is too small to fully confine the nucleus to a unique shape under the cortex. In this case, with a large excess nuclear surface area, given cell volume and spread radius, the nuclear and cortical surfaces would instead be expected to be separated and the nucleus would take on any the infinite possible nuclear shapes (with variable curvature and *τ*_nuc_ = 0) confined under a spherical cap-shape cell cortex.

**Figure 5.**
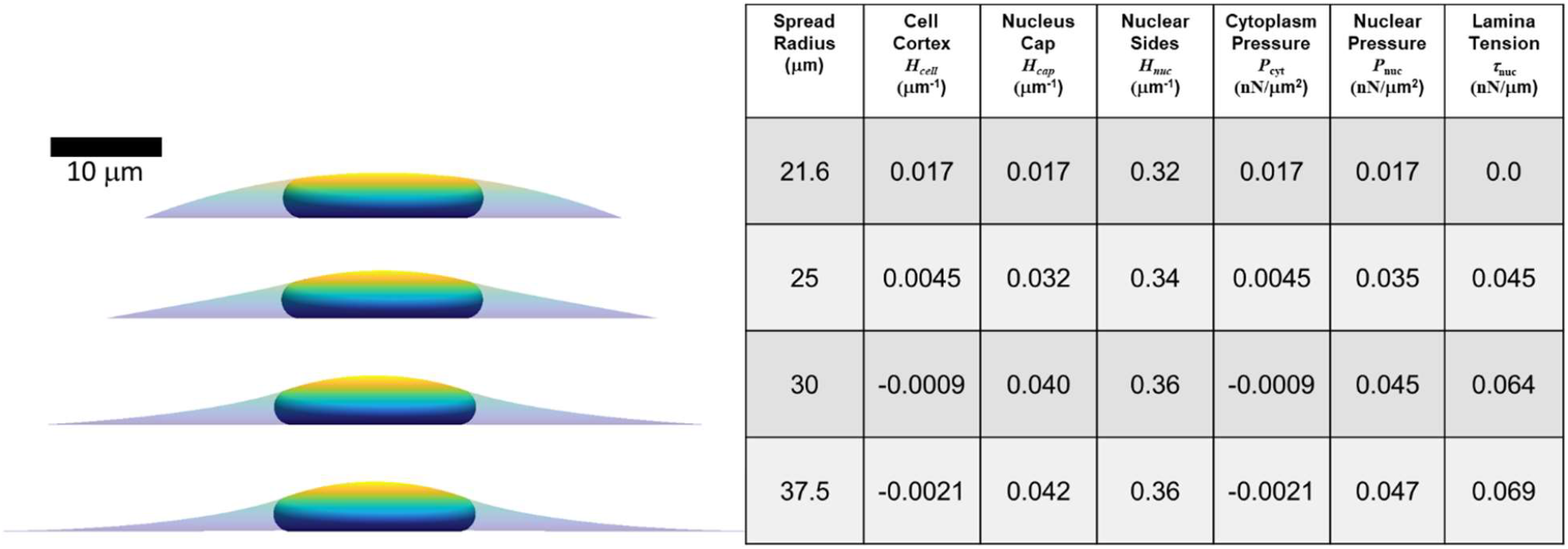
Calculated axisymmetric nuclear shapes, curvatures, and pressures, for cells with varying spread radius, 40% excess lamina area, and the same parameters used in Figure 4 (*V =* 900 µm^3^, *V*_cell_ = 3.4*V, τ*_cell_ = 0.5 nN/µm). The spread radius ranges from the minimum (21.6 microns) where the lamina is under positive tension to the maximum radius (37.5 microns) where the cell edge becomes parallel to the substratum.

### 3.2 Nuclear shapes in cells in an epithelial monolayer or isolated in a well

The shapes of epithelial cells in a monolayer *in vivo* can vary from columnar to cuboidal to squamous. We have previously reported that MCF10A breast epithelial cells in culture monolayers exhibit a flattened morphology with disk-shaped nuclei and remarkably uniform nuclear heights (Figure 6A; [18]). While pulling or compressive stress transmitted to the nucleus from the moving cell boundaries is likely involved in shaping the nucleus during changes in cell shape due to viscous forces [20; 34], maintaining the resulting disk-like nuclear shape does not necessarily require cellular forces and can arise entirely from the constraints on excess area and cell volume, as indicated in Figure 2B. This nuclear shape along with the cell shape can also be predicted for a given cell spreading area, as shown in Figure 6A, using the approach in section 3.1, but now with the vertical position of the cell edge also specified. Interestingly, there is a unique cell radius that yields a flat apical surface of the nucleus; any larger radius would instead have concave regions of lower cell height between the adjacent nuclei in the monolayer, which was not observed experimentally. Nuclei were similarly flat and disk-like when isolated cells were cultured within a 5-µm deep well [18], though in this case the cell cortex was not flat; rather, it exhibited a concave shape with a meniscus due to cell spreading up the side walls (Figure 6B, [18]). These shapes of the cell and nucleus can also be predicted from the model by constraining both the radial and vertical edge position in an axisymmetric well of the same area as used in the experiments. Here, the vertical position of the cell boundary was varied until the observed disk-like nuclear shape was achieved. Importantly, a concave, upwardly curved tensed cortex would not be able to exert a downward compressive force, supporting the model prediction that the disk-like nuclear shape can arise without a downward compressive force on the nucleus. This conclusion holds for either the cell in the monolayer or the cell in the well, because the apical nuclear surface appears flat (*H*_*cap*_ = 0) in both cases. From Eq. 5, a flat apical cap implies that nuclear pressure is zero relative to the cell surroundings, but the lamina tension is predicted to be positive, and the cytoplasmic pressure is negative, for the cell in the well, due to the negative value of *H*_*cell*_. In contrast, the lamina tension and the cytoplasmic pressure for the monolayer cell are predicted be zero since both *H*_*cell*_ and *H*_*cap*_ are near zero. Since there is nothing evident in these experiments that would prevent cells from spreading further (making *H*_*cap*_ > 0, like in typical isolated cells spread on flat substrata), it remains to be explained why the cell spreading is limited in the monolayer or in a well to just reach the point where nuclear pressure is zero.

**Figure 6.**
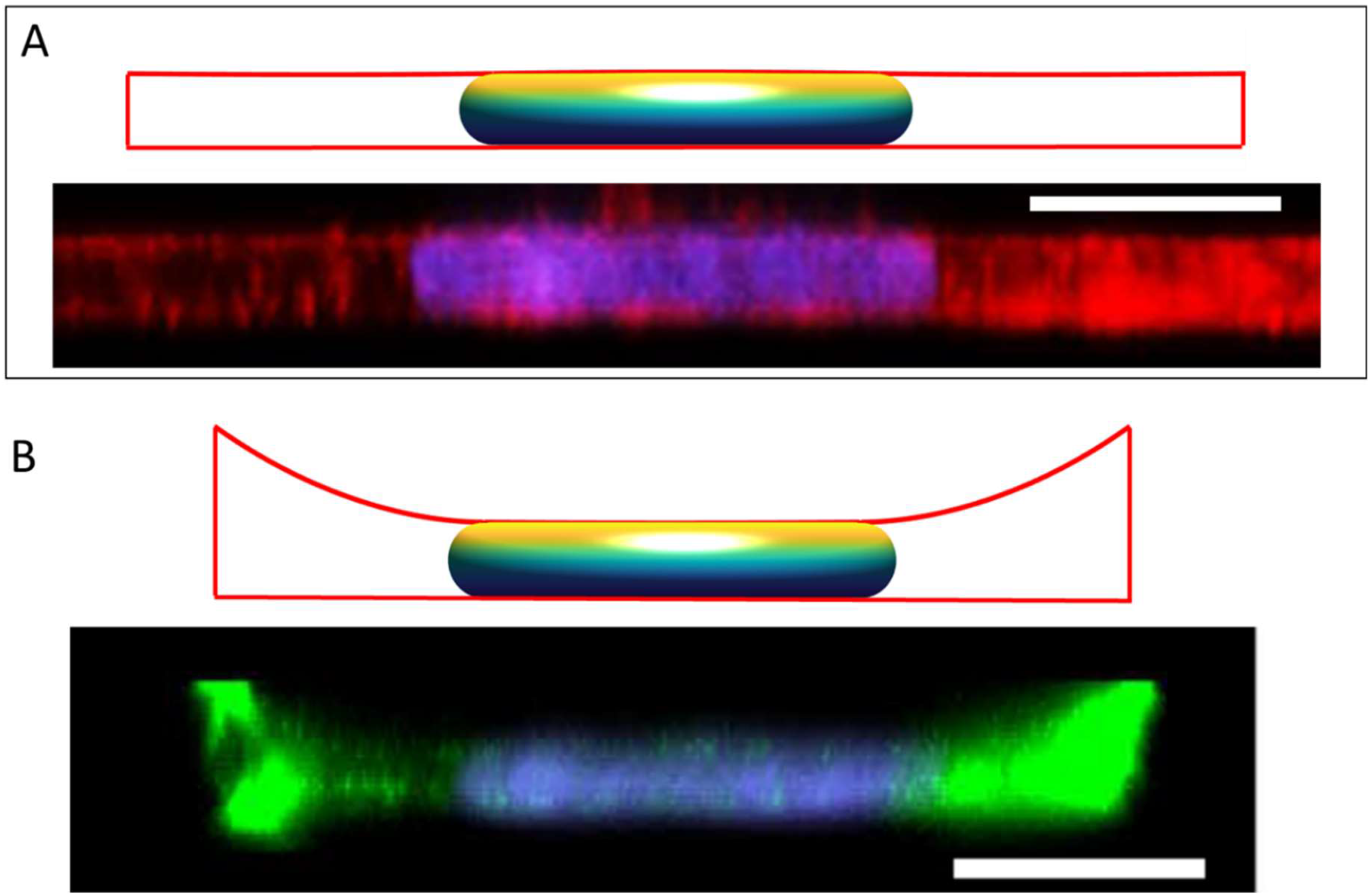
Calculated cell and nuclear shapes for cells in (A) epithelial monolayer; and (B) isolated within well, compared to the experimental *x-z* MCF10A cell and nuclear profiles reported in Neelam et al. *Sci Rep* 2016 [18]. Experimental x-z MCF10A cell profiles were stained for F-actin with Alexa Fluor 488 phalloidin and the nucleus with Hoechst 33342. Calculated cell boundaries are indicated by red lines, with lateral and basal boundaries are imposed by the solid boundaries, and apical surfaces being calculated surfaces of constant mean curvature). The flat disk-shaped nuclei like those observed in epithelial cells are solutions to the axisymmetric model for the cell and nuclear volumes reported in (*V* = 700 µm^3,^ *V*_cell_ = 6.4*V*), and assuming an excess area ε ∼ 65%. The flat apical surface implies that the nuclear pressure relative to the surroundings is nearly zero (see Eq. 5). In the monolayer, the apical cortical surface is also flat, predicting in nearly zero cytoplasmic pressure. But for the cell in the well, the cortex is negatively curved resulting in a negative cytoplasmic pressure and a positive lamina tension. Scale bars are 10 microns.

### 3.3 Nuclear shapes in cells spread on patterned rectangles and lines

Similar to the flat nuclei in flat (spread) cells, elongated nuclear shapes are observed in elongated cells, Versaeval et al. [19] found that nuclear shapes were increasing elongated in cells cultured on rectangles with increasing aspect ratios. To test whether such non-axisymmetric 3D nuclear shapes can be predicted from the cell geometry and the excess nuclear surface area alone, we computed the 3D shapes numerically by minimizing the area of a 3D triangular surface meshes representing the cell cortex and nuclear surface for a given cellular adhesion footprint, under the constraints of constant cell volume, nuclear volume, and nuclear surface area (see Methods). As shown in Figure 7, the computed nuclear shapes closely mirror those reported in [19], assuming an excess area of ∼50%. However, the nuclear volumes required to calculate nuclear sizes consistent with the images and reported aspect ratios were roughly two times larger than the volumes reported in [19], which were calculated assuming ellipsoid nuclear shapes. This difference is likely due to the fact that that the ellipsoid approximation significantly underestimates the volume in the z-direction for a nucleus that is lies nearly flat on the apical and basal surfaces. Like the nuclear shapes in [19], the nuclei are predicted to be much flatter vertically than horizontally, with oval x-y profiles with aspect ratios that mirror the cellular aspect ratios. Importantly, these shapes are again explained purely from the geometric constraints irrespective of the magnitude of the cellular forces, and do not require invoking an elastic force balance like that proposed in [19].

**Figure 7.**
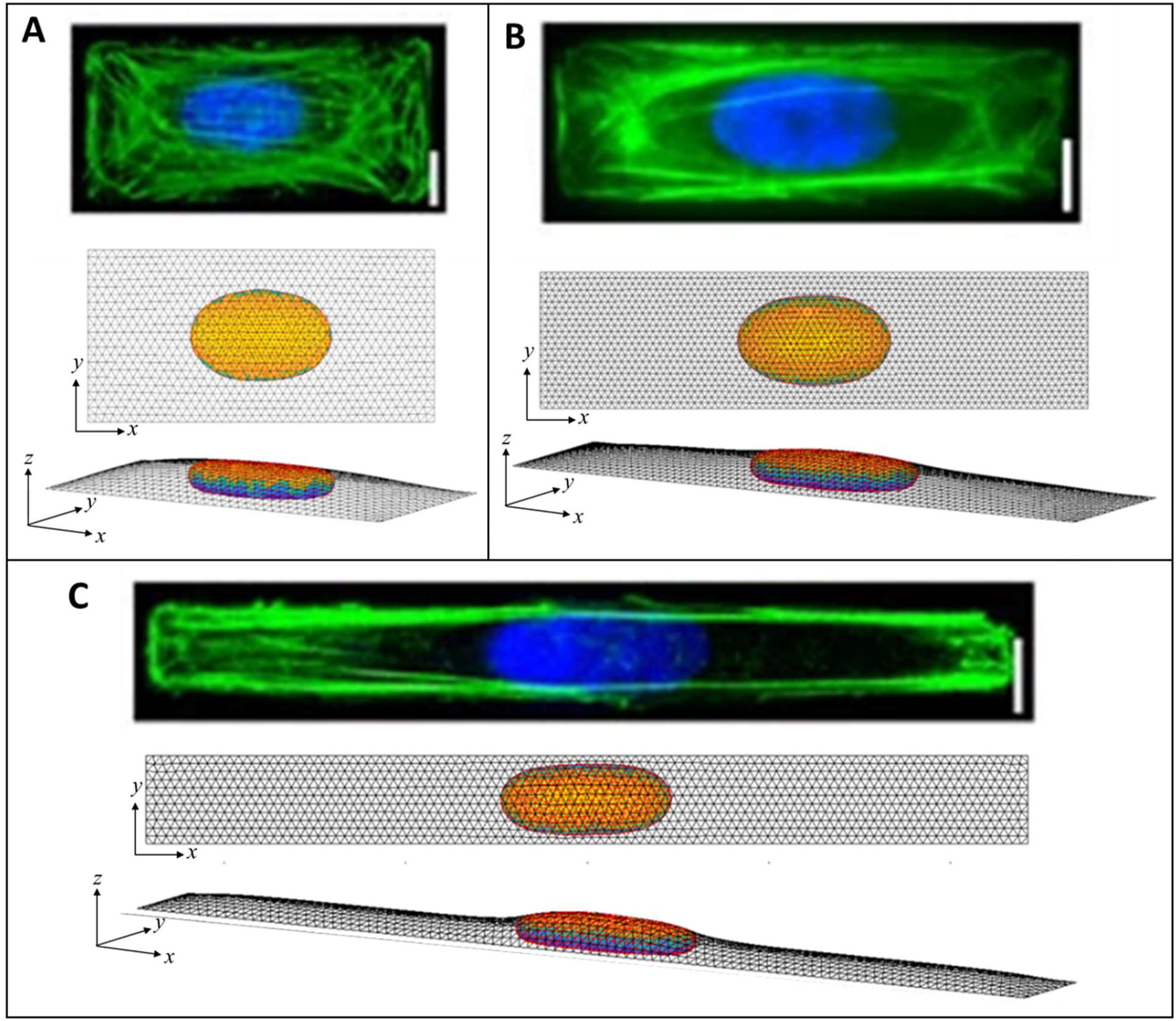
Calculated 3D cell and nuclear shapes compared to data of primary human umbilical vein endothelial cells spread on patterned 1600-μm^2^ rectangles of varying aspect ratio, as reported by Versaeval et al. *Nat Comm* 2012 [19]. Calculated *x-z* nuclear profiles and nuclear heights closely agree with the experimental nuclear shapes, with similarly increasing nuclear lengths with aspect ratios of (A) 2:1, (B) 4:1, and (C) 10:1. Scale bars are 10 microns.

### 3.4 Nuclear shapes with deep indentations

Lastly, we test the ability of the geometric model to explain the overall nuclear shapes when the nucleus develops deep nuclear invaginations, as we have recently reported in nuclei impinging against microposts [43]. In these experiments, nuclei in migrating cells contact the microposts, creating invaginations into the nuclear lamina (Figure 8), similar to the deformation of a liquid drop with surface tension. Here we calculated remarkably similar nuclear shapes by translating a 1-micron diameter micropost toward the nuclear center while recursively calculating the deformed equilibrium nuclear shape. We did not otherwise model the interaction between the micropost and cell, consistent with the experimental observation that the micropost was fully engulfed in the cytoplasm [43]. The calculated nuclear shapes for *ε* = 40-60% closely resemble the experimentally observed nuclear shapes (Figure 8). Moreover, assuming a cortical tension of ∼0.5 nN/um [42], the force on the post was found to be ∼ 0.3-1 nN for *ε* = 40-60%, by accounting for the lamina surface tension from Eq. 5 and the length of interaction with micropost. This is close experimental values of 1-2 nN reported in [43]. These computational findings are consistent with the interpretation of the shapes in [43], that the invaginations reflect the lamina surface tension surrounding a pressurized yet compliant nuclear interior, rather than an elastic deformation of the nucleus. The excess surface area permits such extreme shape changes again without stretching the lamina or compressing the nuclear volume.

**Figure 8.**
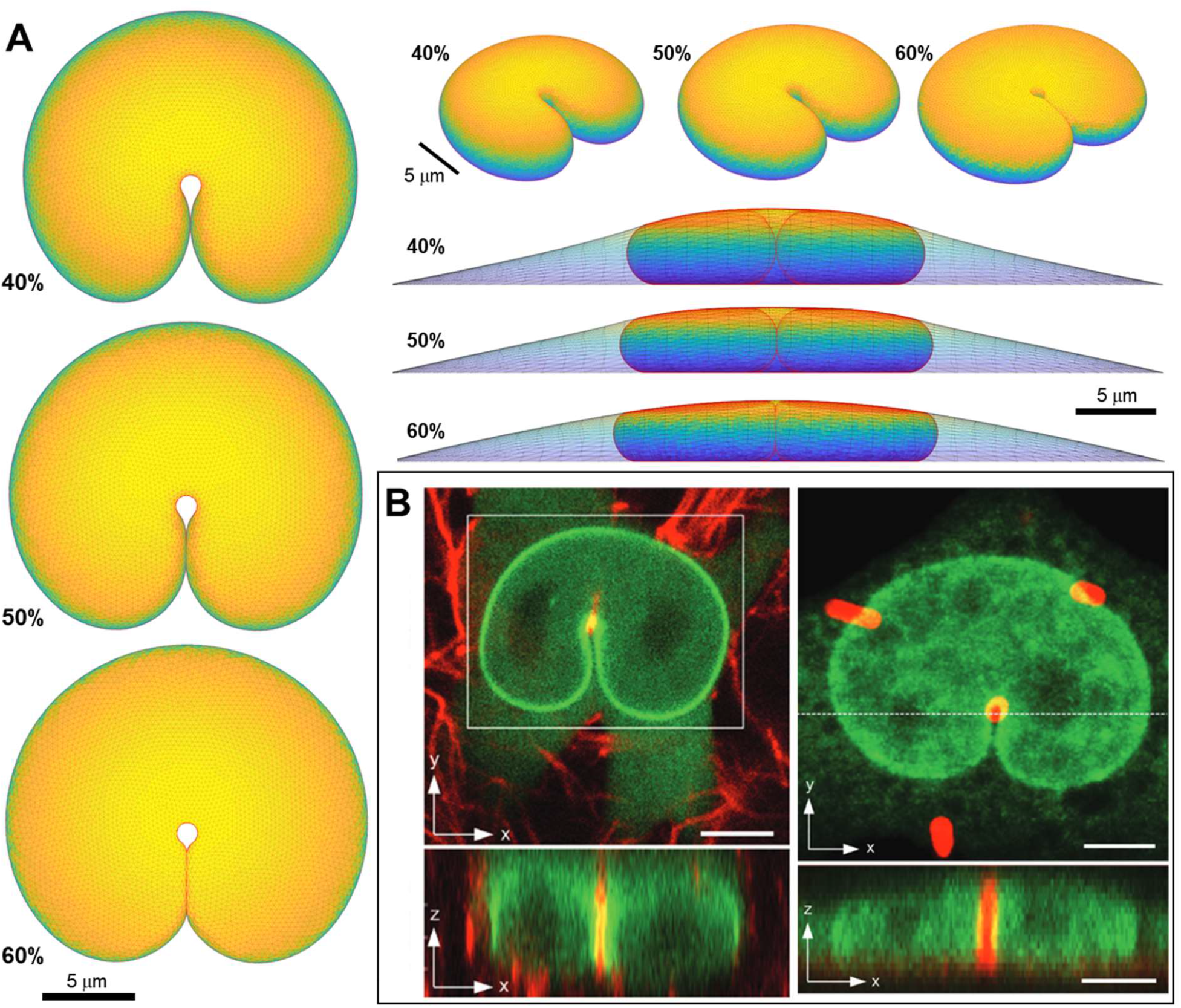
Predicted 3D shape from geometric model for nucleus deformed by a 1-um diameter micropost compared to 3D imaging in Katiyar et al. *Adv Sci* 2022 [44]. (A) Calculated shapes from three different viewing angles (top, elevated, and side with cell shape also shown) are for nuclear volume, *V =* 900 μm^3^, cell volume, *V*_cell_ = 3.4*V*, and three different fractional excess areas. 40%, 50%, and 60%, (B) Experimental shapes for nuclei indented by (i) collagen fiber and (ii) one micron micropost (in red). Green is GFP-lamin (scale bar is 5 microns).

## 4. Discussion

An oval or circular nucleus with a smooth, curved contour is a ubiquitous, striking feature of a cultured eukaryotic cell. Yet, an explanation for its smooth appearance has remained elusive. In contrast to our geometric explanation, deformed non-spherical nuclear shapes have been widely assumed to result from a balance of cytoskeletal forces on the elastically deformed nucleus (reviewed in [34] and [1]), with the resting state of the nucleus assumed to be an undeformed sphere. Our results contradict this notion, showing that geometric considerations alone can parsimoniously explain a wide range of nuclear shapes observed in different experimental contexts, independent of the magnitude of cytoskeletal forces. The nuclear shape calculations require no parameters other than the (excess) nuclear surface area and the cell and nuclear volumes. Before the limiting shapes are reached, excess lamina surface area is predicted to allow the nucleus to undergo dynamic shape deformations at constant surface area and constant volume. These deformations can occur because they do not require areal expansion of the lamina or compression of the nuclear volume. This principle explains why the shape changes of the nucleus conform to cellular shape changes during cell spreading [20] and cell crawling [34; 38], before reaching the limiting shapes modeled here. In these dynamic situations, cytoskeletal forces and cytoskeletal linkages to the nucleus involved in transmitting viscous or viscoelastic stresses to the nucleus are expected to be important in driving the nucleus to the limiting shapes. That is, the pathway and time required to reach the limiting shapes likely depends on the nature, magnitude, and transmission of the cytoskeletal forces, while the ultimate limiting shape is geometrically determined for a given cell shape. For example, disruption of the LINC complex, which connects the cytoskeleton to the nucleus, slows nuclear flattening during cell spreading but does not affect the ultimate limiting shape [20].

It is surprising that the model can so effectively capture nuclear shapes in various cell geometries with so few parameters despite several simplifying assumptions. The assumption of surfaces of constant mean curvature implicitly assumes that the nuclear pressure *P*_*nuc*_ and cytoplasmic pressure *P*_*cyt*_ are spatially unform and isotropic (the latter term accounting for both the hydrostatic pressure in the cytosol and contractile tension in cytoskeletal network phase [20]). Constant mean curvature also implies that the surface tensions in the cell cortex (*τ*_*cell*_) and the nuclear lamina (*τ*_*nuc*_) are spatially uniform and isotropic. It is likely that anisotropic surface tension may impact the directionality of cell surface curvature, especially in highly elongated cells where stress fibers in the cell cortex tend to align with the cell’s long axis, but such anisotropy still does not appear to be a primary driver for nuclear shape. The model also ignores the effect of cytoskeletal structures (e.g., organelles and stress fibers) pressing against the nucleus. However, these do not appear to greatly change the smooth nuclear shape when the lamina is tensed, and their impact on the nuclear shape is likely to be more pronounced in less spread cells where the lamina is not tensed.

The key model assumption that the lamina has excess surface area is clearly evident in the folds, wrinkles, and surface undulations seen in 3D images of rounded nuclei, falsifying the notion that the resting state of the lamina is spherical (Figure 1). Moreover, our previous work has shown that removal of cytoskeletal forces does not cause relaxation of the nucleus to a spherical morphology, implying nuclear deformations in spread cells are irreversible [38]. Consistent with these findings, elastic forces in the nuclear interior have been found to rapidly dissipate on the time scale of seconds and that the nuclear contents behave as a viscous fluid on this time scale [45; 46]. From literature estimates of the area dilation modulus (∼390 nN/µm; [47]), the nuclear bulk modulus (∼5 nN/µm^2^ [48]), and the cortical tension (∼0.5 nN/µm; [42]), the areal extension of the lamina and changes in volume due to compression in the flattened nucleus during spreading are expected to be less than 1%, based on pressures and tensions calculated from Eq. 5. Even if some nuclear compression or areal expansion were to occur when the lamina becomes smoothed and tensed, this would not weigh against the key conclusions that shape is primarily limited by the geometric constraints of lamina area and nuclear volume, and that the excess area permits a wide range of shapes at constant volume and lamina area before the limiting shape is reached. The modeling approach can easily accommodate nuclear compressibility and lamina-areal expansion by incorporating a bulk modulus and area modulus, as we have done previously in Li et al. [20].

Because the calculated nuclear shapes are unique, limiting geometric shapes constrained by constant lamina surface area and constant nuclear volume, any resistance to bending of the nuclear lamina and/or the nuclear envelope is not relevant for predicting the shapes. If bending stiffness of the lamina is large enough, it could, in principle, affect the calculated pressures and tensions. The surface bending energy per unit area can be calculated as *E*_*bend*_ = 2*k*_*c*_*H*^2^ [49], where *k* is the bending modulus and *H* is the mean curvature. For *k*_*c*_ ∼ 0.4 nN-µm [50; 51; 52], *E*_*bend*_ < 10^−4^ nN-µm using *H*∼0.4 µm^−1^ for the most-curved nuclear surface regions. By comparison, the estimated surface tension in the lamina is *τ*_*nuc*_ ∼ 5×10^−2^ nN-µm (Figure 4 and 5). Thus, the bending energy of the lamina, even in the regions of highest curvature, should be negligible, being 2-3 orders of magnitude smaller than the calculated values of lamina surface tension. Furthermore, the nuclear lamina commonly exhibits folds and wrinkles on time scales of tens of minutes or longer during cell spreading before a limiting shape with a smoothed lamina is reached [44]. Thus, it is unlikely that resistance to bending of the lamina or the other envelope components plays a significant role in driving nuclear shape changes, at least on the longer time and length scales considered here.

Our results predict that nuclear pressure and lamina surface tension should arise in cell geometries that fully unfold the lamina excess area (which is the case for most mammalian cells in culture). This emphasizes the key mechanical role that the nuclear lamina plays in imparting surface tension to the nuclear surface. The lamina can protect the nucleus from extreme deformations, while otherwise permitting mechanical compliance due to its excess area. For example, nuclear pressure and the resulting lamina tension explain the source and magnitude of forces exerted on microposts which indent nuclei [44] in migrating cells (c.f. Figure 8). They also explain how nuclei pass by the obstacles unimpeded. In contrast, cells without lamin A/C appeared to lack surface tension, resulting in the entanglement of highly deformed nuclei on the obstacles. Similarly, nuclei in lamin A/C-null cells flatten more than in wildtype spread cells [20], again implying that the constant-surface area constraint on nuclear shapes requires lamin A/C.

The prediction of tension in the lamina upon unfolding is also consistent with the observation that mechanosensitive yes-associated protein (YAP) import to the nucleus correlates with nuclear unwrinkling in cells in 2D culture [37], and that nuclei rupture during stretching of the lamina during cell migration through confining spaces[53]. As YAP translocation regulates gene expression, while deformation and rupture can promote DNA damage and tumorigenesis, the mechanical state of the nuclear lamina predicted by the model is likely to be important in both healthy and diseased cells. Abnormal nuclear morphologies in cultured diseased cells (such as tumor cells or progeric cells) may also be in part due to changes in nuclear pressure and tension in the lamina, which is a possibility worth exploring in the future.

In summary, the simple geometric principle invoking excess surface area explains how large nuclear deformations seen in various cell geometries do not require that the nucleus be subjected to a large force. Rather, limiting deformed nuclear shapes are geometrically determined and not mechanically determined. We anticipate that future application of this principle will yield further fundamental insights into the relationships between force, nuclear deformation, and cell function in healthy and diseased tissues.

## 5. Methods

### 5.1 Geometric prediction of cell and nuclear shapes in axisymmetric spread cells

Here we derive the equations describing the general shape, surface area, and volume of an axisymmetric surface of constant mean curvature. Next, the equations are used for the various cell surfaces to predict the axisymmetric cell shape and nuclear shape. Let *z* be the vertical height of an axisymmetric surface and *R* be the radial distance from the axis of symmetry. Let *θ* be the angle between the vertical z-direction and the tangent to the interface, such that

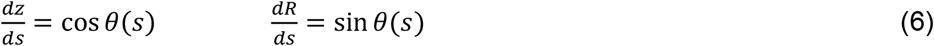

where *s* is distance along the arc-length. Accounting for the curvatures along the arc-length and the orthogonal azimuthal direction, the mean curvature of the axisymmetric surface is

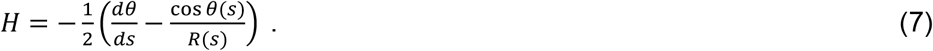

Upon changing variables to *R* and *z*, Eq 7 can be written,

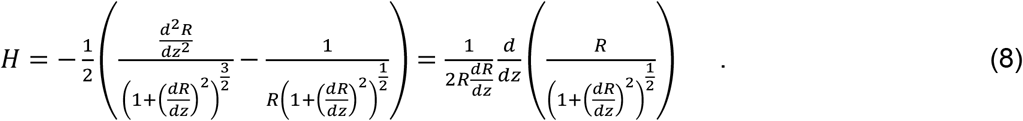

Rearranging and integrating with respect to *z* yields

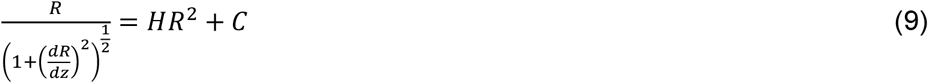

where *C* is a constant, thus

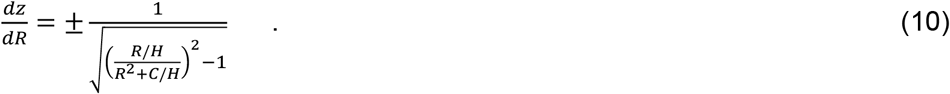

To integrate Eq. 10, it can be rewritten by replacing parameters *C* and *H* with new parameters *α* and *β*, such that

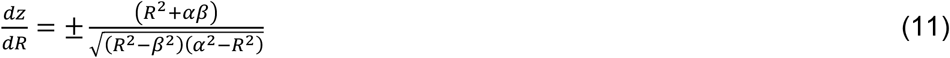

where

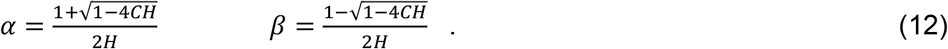

Note that the original parameters *C* and *H* can be recovered from

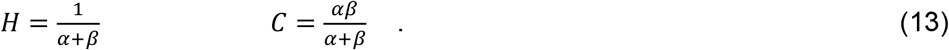

Now, a new variable *ϕ* is introduced to replace *R* such that

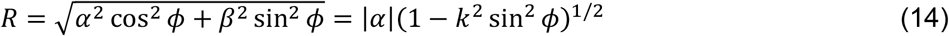

Where

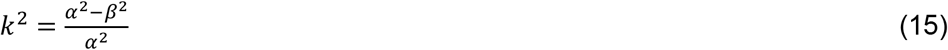

such that

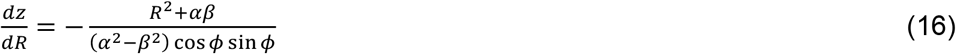

(both sign possibilities in Eq. 11 are now accounted for by allowing positive and negative values of *ϕ*). Because

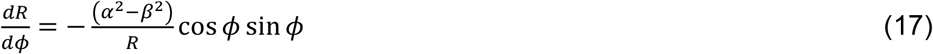

this change in variable simplifies Eq. 6 to be:

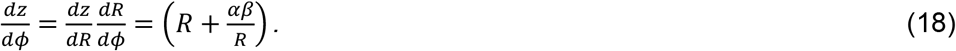

Note that *ds*^2^ = (*β* + *α*)^2^*dϕ*^2^ = *H*^2^*dϕ*^2^, providing a physical interpretation of *ϕ* as the arc length scaled by the mean curvature. Integrating Eq. 8 from initial value *ϕ*_0_ provides

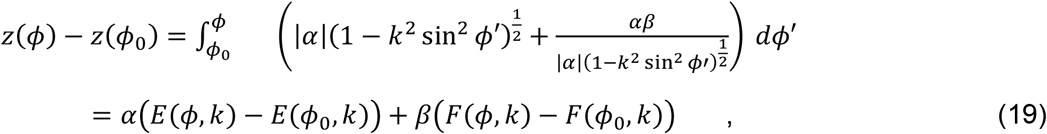

where and *F*(*ϕ, k*) and *E*(*ϕ, k*) are incomplete elliptical integrals of the first and second kinds, respectively, and

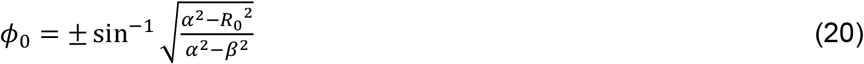

The negative sign in Eq. 20 is used when 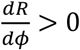 at *R*_0_ ≡ *R*(*ϕ*_0_), noting Eq. 17. The surface area is obtained by integrating the arc length rotated around the z-axis:

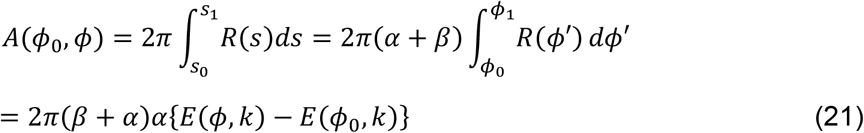

Finally, the enclosed volume between *z*_0_ and *z* is similarly obtained:

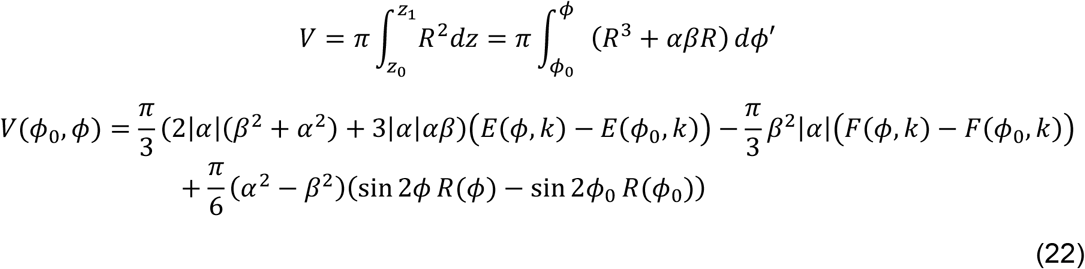

The above results will now be applied to cortical, nuclear, and joint surfaces, while accounting for the boundary conditions, constraints, and the junction between them at point (*R*_1_, *z*_1_) (see Figure 9). First, the cortex surface (*R*(*ϕ*), *z*(*ϕ*)) and mean curvature *H*_*cell*_ is calculated between (*R*_1_, *z*_1_) and the cell edge at (*R*_0_, *z*_0_). In this case,

**Figure 9.**
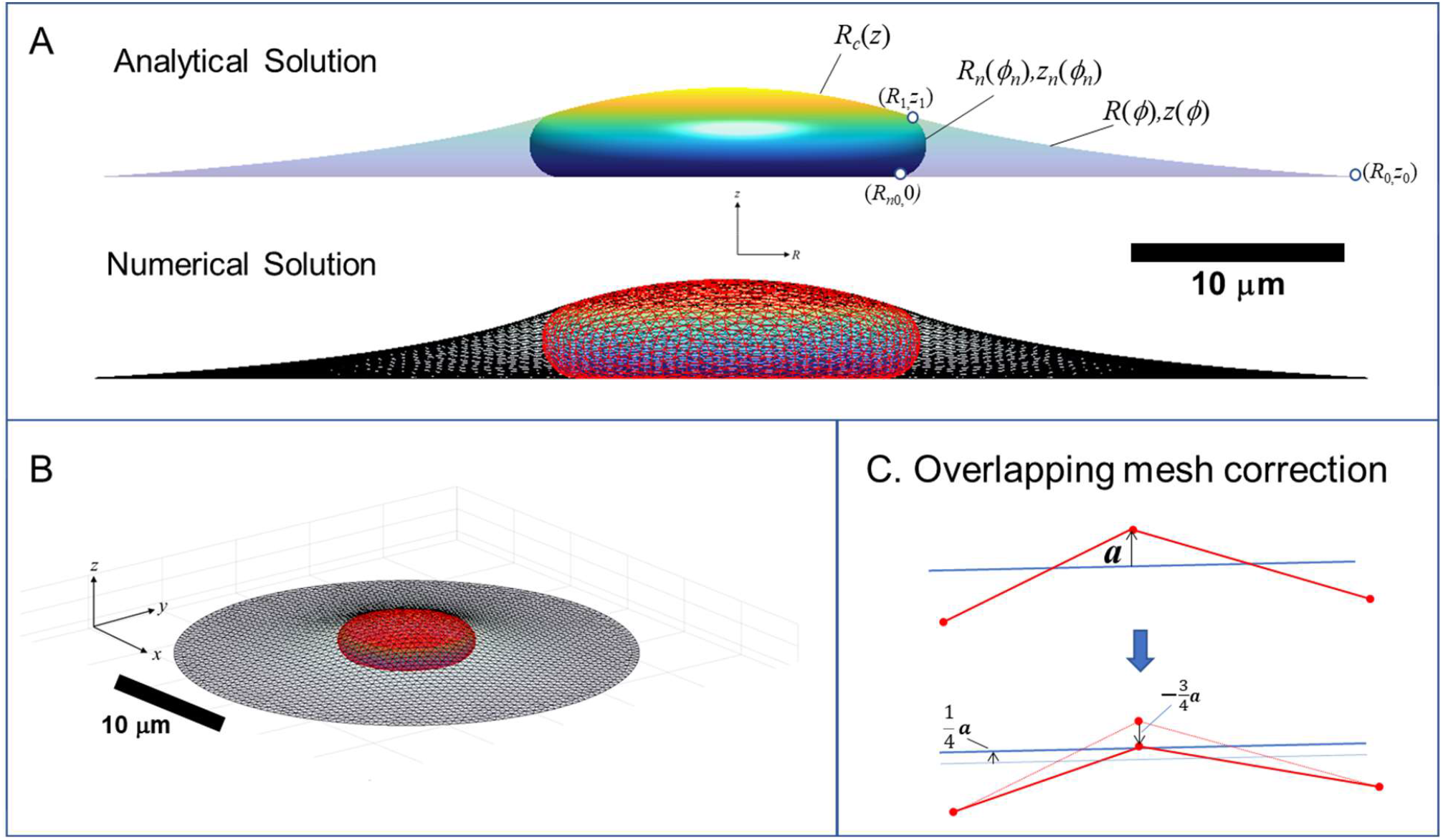
Calculation of three-dimensional cell and nuclear shapes. (A) Analytical axisymmetric surfaces of constant mean curvature for the cortical surface interface with the surroundings, (*R,z*), nucleus-cytoplasm interface containing the nuclear lamina, (*R*_*n*_,*z*_*n*_), and joint nucleus-cortical interface with the surroundings, *R*_*c*_(*z*). Surfaces are matched at point (*R*_1_,*z*_1_), and *R*_*n*_ is the radius to which the nucleus presses against the substratum. The calculation shown here is for *V =* 900 µm^3^, *V*_cell_ = 3.4*V, ε* = 0.45, and spread radius of *R*_0_ = 30 µm (*z*_0_ = 0), (B) 3D numerical calculation for the same conditions obtained minimizing the surface areas and simultaneously optimizing the triangular mesh by maintaining a centroidal Voronoi tessellation using the algorithm in [54]. Nuclear and cortical surfaces were solved simultaneously for the for the given adhesion footprint under the constraints of constant nuclear surface area, nuclear volume, and cell volume. The method is validated by close agreement with the exact analytical solution for an axisymmetric spread cell. (B) Elevated perspective of the same 3D cell from (A). (C) Illustration of algorithm for preventing the crossing of meshes. When a vertex point (shown as red dot on projected triangle edges) crosses an opposing mesh triangle surface (indicated by blue line), with distance vector from the nearest triangle surface ***a***, the vertex is pushed back by distance 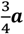, and each of the three vertices of the opposing triangle is pushed forward by 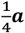. In this way, the vertex ends up on the plane of the triangle, and the forces balance on the two surfaces, with each vertex of the opposing triangle shares and equal share of the force.

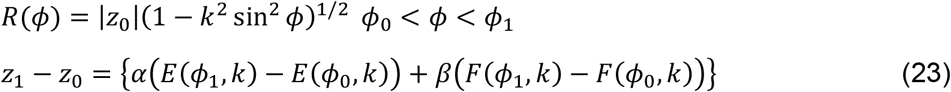

where

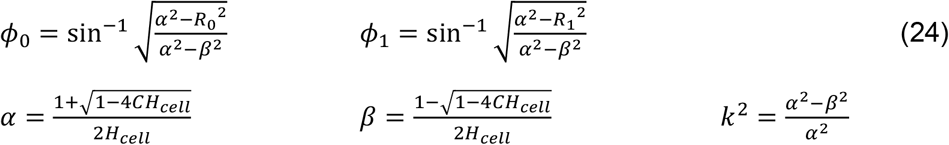

with parameter *C* to be determined from the boundary conditions below. Similarly, the nuclear surface profile (*R*_*n*_(*ϕ*_*n*_), *z*_*n*_(*ϕ*_*n*_)) between the contact point on the substratum, *R*_*n*0_ and *R*_1_is

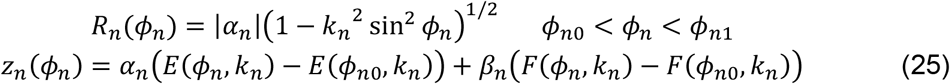

\where, for curvature *H*_*nuc*_ and parameter *C*_*n*_ from Eq. 5, we have

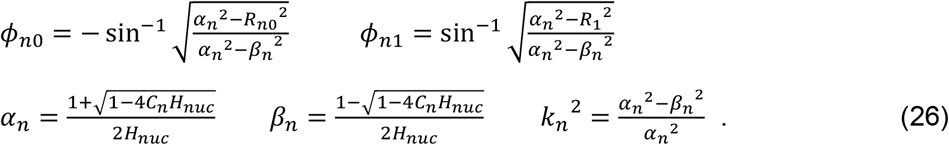

Finally, the joint cortex-lamina surface capping the cell, *R*_*c*_(*z*), is derived from *R*_1_ to the cell apex where *R*_*c*_ = 0. From Eq. 9, the constant *C* must be zero, such that the shape takes the form of a spherical cap of radius *H*_*cap*_ ^−1^ and height

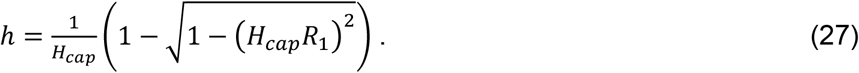

The volume and surface area of a spherical cap of height are *h* and radius *H*_*cap*_ ^−1^ are

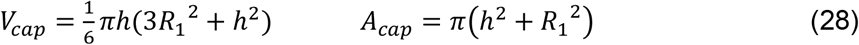

The parameters *C*_*c*_ *C*_*n*_ *R*_*n*0_ can now be obtained by considering the boundary conditions. Assuming no attachment between the nuclear envelope and the cortex at the junction at *R*_1_ to sustain a normal force, the lamina slope is continuous and the positions and tangents of the three surfaces must be equal at (*R*_1_, *z*_1_), i.e.

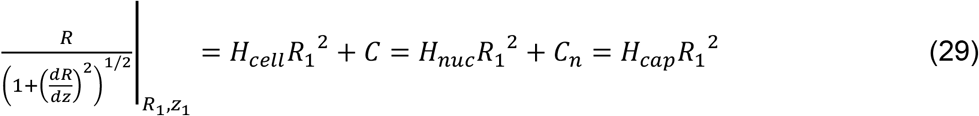

thus

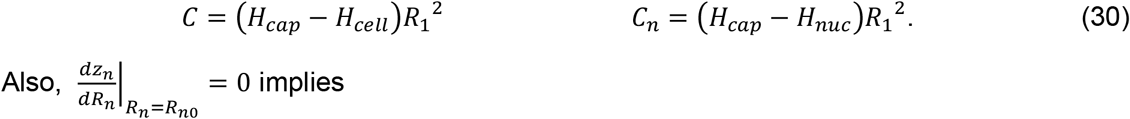

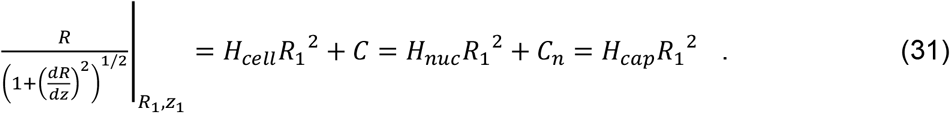

*H*_*nuc*_ *R*_*n*0_^2^ + *C*_*n*_ = 0, such that

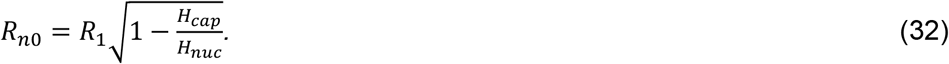

At this point, the cortex and nuclear profiles can be calculated for given values of parameters *R*_1_, *H*_*cell*_, *H*_*cap*_ and *H*_*nuc*_, and attachment-point boundary condition, (*R*_0_, *z*_0_). These values are determined implicitly by applying four constraints, namely setting *z*_*n*_ *ϕ*_*n*,1_ = *z*_1_, and by fixing the nuclear surface area and cell and nuclear volumes

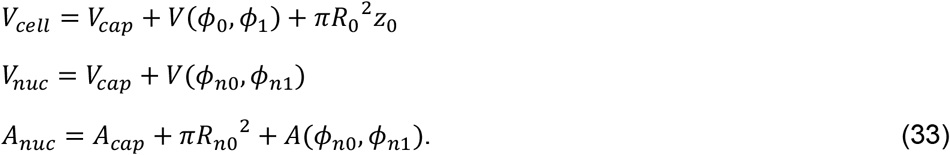

Here, Eq. 21 and 22 are used to calculate *A*(*ϕ*_*n*0_, *ϕ*_*n*1_), *V*(*ϕ*_0_, *ϕ*_1_), and *V*(*ϕ*_*n*0_, *ϕ*_*n*1_), and Eq. 28 is used for *V*_*cap*_ and *A*_*cap*_. It should be noted that some ranges of (*R*_0_, *z*_0_) permit the solutions with *H*_*cell*_ > *H*_*cap*_, which would not be physically possible (and is not generally observed) since it implies negative surface tension on either the cortex or lamina. For cell spreading on a substratum, i.e., *z*_0_ = 0, there is a minimum radius *R*_0_ where *H*_*cell*_ = *H*_*cap*_, at which point the lamina becomes under tension due to compression from the cortex (see Figure 3). Below this radius, the cortex need not impinge on the nucleus and the nucleus can take a wide range of possible shapes that satisfy the constant volume and area constraints.

### 4.2 Calculation of cell and nuclear shape in 3D geometries

Unlike the axisymmetric case, three dimensional geometries do not generally permit analytical solutions of cell and nuclear shapes. Instead, surfaces of constant mean curvature for the cell and nucleus were calculated using an optimization algorithm to minimize the surface areas under the constraints of constant cell and nuclear volume for a given cell geometry. Surfaces of constant mean curvature with constraints were generated using the approach of Pan et al. [54] which minimizes surface area of a triangular mesh *ℳ*(*X*) (with *N* vertices at positions, 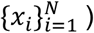 while simultaneously maintaining a centroidal Voronoi surface tessellation (with Voronoi cells, 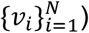). In this approach, a surface is constant curvature is achieved by optimizing the tessellation based on the following energy function of *X*:

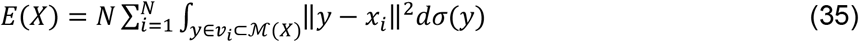

where the integrals are over the area of the Voronoi cells surrounding each of the vertices. Accounting for the volume and area constraints, *A*(*X*_*nuc*_) = *A*_*nuc*_ and *V*(*X*_*nuc*_) = *V*_*nuc*_, the final cell and nuclear vertex positions, *X*_*cell*_ and *X*_*nuc*_, respectively, were obtained by minimizing the total energy function,

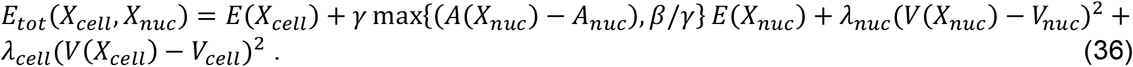

The area and volume stiffness parameters, *γ, λ*_*nuc*_, and *λ*_*ncell*_ were progressively increased toward large values (>10^5^) after each shape convergence until |*V*(*X*_*nuc*_) − *V*_*nuc*_| /*V*_*nuc*_, |*A*(*X*_*nuc*_) − *A*_*nuc*_|/*A*_*nuc*_, and |*V*(*X*_*cell*_) − *V*_*cell*_|/*V*_*cell*_ are all less than 10^−4^. The central term ensures that a small background surface tension (reflected by small value of parameter *β*) is assigned during the optimization when *A*(*X*_*nuc*_) < *A*_*nuc*_ to maintain a smooth mesh without wrinkles and buckles during the optimization. Boundary conditions at the edges of the cell adhesion area were imposed by fixing vertex positions on edge of the cell adhesion area, and the Voronoi tessellation was maintained by flipping edges when two opposite angles of two adjacent triangles summed to be greater than π (see [54] for algorithm details on flipping edges). The initial mesh of nearly equilateral triangles was generated using the DISTMESH algorithm [55]. A steepest descent optimization algorithm was used to converge to an equilibrium shape with surfaces of constant curvature under the area and volume constraints. The algorithm was validated by comparison to analytical solutions of the axisymmetric case (Figure 9A,B).

In the optimization algorithm, opposing mesh surfaces were prevented from overlapping by correcting displacements of vertices which cross the triangle faces of the opposing mesh surface, while simultaneously displacing the vertices of the opposing triangle in the opposite direction (Figure 9C). If ***a*** is the vector orthogonal from the surface to the uncorrected overlapping position of the vertex point, then the vertex position correction is 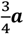 for the vertex, and each of the three vertices of the opposing triangle was adjusted by 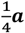. In this way, the corrected vertex position ends in the plane of the opposing triangle, the surface forces remain equal and opposite. Similarly, vertex positions that cross solid surfaces (i.e., the substratum or the micropost) were adjusted to the projected position on the solid surface.

## ACKNOWLEDGEMENTS

This work was supported by NIH U01 CA225566 (T.P.L. and R.B.D.), and a CPRIT established investigator award Grant No. RR200043 (T.P.L.). We thank Christina Dubell for collecting confocal images of the nucleus and cell in some of the figures.

## References

[1] T.P. Lele, R.B. Dickinson, and G.G. Gundersen, Mechanical principles of nuclear shaping and positioning. J Cell Biol 217 (2018) 3330–3342.

[2] T. Harada, J. Swift, J. Irianto, J.W. Shin, K.R. Spinler, A. Athirasala, R. Diegmiller, P.C. Dingal, I.L. Ivanovska, and D.E. Discher, Nuclear lamin stiffness is a barrier to 3D migration, but softness can limit survival. J Cell Biol 204 (2014) 669–82.

[3] E.R. Smith, Y. Meng, R. Moore, J.D. Tse, A.G. Xu, and X.X. Xu, Nuclear envelope structural proteins facilitate nuclear shape changes accompanying embryonic differentiation and fidelity of gene expression. BMC Cell Biol 18 (2017) 8.

[4] Y.K. Wu, H. Umeshima, J. Kurisu, and M. Kengaku, Nesprins and opposing microtubule motors generate a point force that drives directional nuclear motion in migrating neurons. Development 145 (2018).

[5] N.Y. Chen, Y. Yang, T.A. Weston, J.N. Belling, P. Heizer, Y. Tu, P. Kim, L. Edillo, S.J. Jonas, P.S. Weiss, L.G. Fong, and S.G. Young, An absence of lamin B1 in migrating neurons causes nuclear membrane ruptures and cell death. Proc Natl Acad Sci U S A 116 (2019) 25870–25879.

[6] A.C. Rowat, D.E. Jaalouk, M. Zwerger, W.L. Ung, I.A. Eydelnant, D.E. Olins, A.L. Olins, H. Herrmann, D.A. Weitz, and J. Lammerding, Nuclear envelope composition determines the ability of neutrophil-type cells to passage through micron-scale constrictions. J Biol Chem 288 (2013) 8610–8618.

[7] D. Lorber, R. Rotkopf, and T. Volk, A minimal constraint device for imaging nuclei in live Drosophila contractile larval muscles reveals novel nuclear mechanical dynamics. Lab Chip 20 (2020) 2100–2112.

[8] A.J. Lomakin, C.J. Cattin, D. Cuvelier, Z. Alraies, M. Molina, G.P.F. Nader, N. Srivastava, P.J. Saez, J.M. Garcia-Arcos, I.Y. Zhitnyak, A. Bhargava, M.K. Driscoll, E.S. Welf, R. Fiolka, R.J. Petrie, N.S. De Silva, J.M. Gonzalez-Granado, N. Manel, A.M. Lennon-Dumenil, D.J. Muller, and M. Piel, The nucleus acts as a ruler tailoring cell responses to spatial constraints. Science 370 (2020).

[9] A. Elosegui-Artola, I. Andreu, A.E.M. Beedle, A. Lezamiz, M. Uroz, A.J. Kosmalska, R. Oria, J.Z. Kechagia, P. Rico-Lastres, A.L. Le Roux, C.M. Shanahan, X. Trepat, D. Navajas, S. Garcia-Manyes, and P. Roca-Cusachs, Force Triggers YAP Nuclear Entry by Regulating Transport across Nuclear Pores. Cell 171 (2017) 1397-1410.e14.

[10] S. Dupont, L. Morsut, M. Aragona, E. Enzo, S. Giulitti, M. Cordenonsi, F. Zanconato, J. Le Digabel, M. Forcato, S. Bicciato, N. Elvassore, and S. Piccolo, Role of YAP/TAZ in mechanotransduction. Nature 474 (2011) 179–83.

[11] Y. Kalukula, A.D. Stephens, J. Lammerding, and S. Gabriele, Mechanics and functional consequences of nuclear deformations. Nat Rev Mol Cell Biol (2022).

[12] C.M. Denais, R.M. Gilbert, P. Isermann, A.L. McGregor, M. te Lindert, B. Weigelin, P.M. Davidson, P. Friedl, K. Wolf, and J. Lammerding, Nuclear envelope rupture and repair during cancer cell migration. Science 352 (2016) 353–8.

[13] G.P.F. Nader, S. Aguera-Gonzalez, F. Routet, M. Gratia, M. Maurin, V. Cancila, C. Cadart, A. Palamidessi, R.N. Ramos, M. San Roman, M. Gentili, A. Yamada, A. Williart, C. Lodillinsky, E. Lagoutte, C. Villard, J.L. Viovy, C. Tripodo, J. Galon, G. Scita, N. Manel, P. Chavrier, and M. Piel, Compromised nuclear envelope integrity drives TREX1-dependent DNA damage and tumor cell invasion. Cell 184 (2021) 5230–5246 e22.

[14] M. Vortmeyer-Krause, M.t. Lindert, J.t. Riet, V.t. Boekhorst, R. Marke, R. Perera, P. Isermann, T. van Oorschot, M. Zwerger, F. Yang, M. Svoreň, A. Madzvamuse, J. Lammerding, P. Friedl, and K. Wolf, Lamin B2 follows lamin A/C-mediated nuclear mechanics and cancer cell invasion efficacy. bioRxiv (2020) 2020.04.07.028969.

[15] K. Wolf, M. Te Lindert, M. Krause, S. Alexander, J. Te Riet, A.L. Willis, R.M. Hoffman, C.G. Figdor, S.J. Weiss, and P. Friedl, Physical limits of cell migration: control by ECM space and nuclear deformation and tuning by proteolysis and traction force. J Cell Biol 201 (2013) 1069–84.

[16] P. Friedl, K. Wolf, and J. Lammerding, Nuclear mechanics during cell migration. Curr Opin Cell Biol 23 (2011) 55–64.

[17] I. Singh, and T.P. Lele, Nuclear morphological abnormalities in cancer – a search for unifying mechanisms, Springer, 2022.

[18] S. Neelam, P. Hayes, Q. Zhang, R. Dickinson, and T. Lele, Vertical uniformity of cells and nuclei in epithelial monolayers. Scientific Reports 6 (2016) 19689.

[19] M. Versaevel, T. Grevesse, and S. Gabriele, Spatial coordination between cell and nuclear shape within micropatterned endothelial cells. Nat Commun 3 (2012) 671.

[20] Y. Li, D. Lovett, Q. Zhang, S. Neelam, R.A. Kuchibhotla, R. Zhu, G.G. Gundersen, T.P. Lele, and R.B. Dickinson, Moving cell boundaries drive nuclear shaping during cell spreading. Biophysical Journal 109 (2015) 670–686.

[21] F. Guilak, J.R. Tedrow, and R. Burgkart, Viscoelastic properties of the cell nucleus. Biochem. Biophys. Res. Commun. 269 (2000) 781–6.

[22] K.N. Dahl, A.J. Engler, J.D. Pajerowski, and D.E. Discher, Power-law rheology of isolated nuclei with deformation mapping of nuclear substructures. Biophys. J. 89 (2005) 2855–64.

[23] J.D. Pajerowski, K.N. Dahl, F.L. Zhong, P.J. Sammak, and D.E. Discher, Physical plasticity of the nucleus in stem cell differentiation. Proc. Natl. Acad. Sci. 104 (2007) 15619–24.

[24] I. Ivanovska, J. Swift, T. Harada, J.D. Pajerowski, and D.E. Discher, Physical plasticity of the nucleus and its manipulation. Methods Cell Biol. 98 (2010) 207–20.

[25] J. Swift, I.L. Ivanovska, A. Buxboim, T. Harada, P.C. Dingal, J. Pinter, J.D. Pajerowski, K.R. Spinler, J.W. Shin, M. Tewari, F. Rehfeldt, D.W. Speicher, and D.E. Discher, Nuclear lamin-A scales with tissue stiffness and enhances matrix-directed differentiation. Science 341 (2013) 1240104.

[26] J.W. Shin, K.R. Spinler, J. Swift, J.A. Chasis, N. Mohandas, and D.E. Discher, Lamins regulate cell trafficking and lineage maturation of adult human hematopoietic cells. Proc. Natl. Acad. Sci. 110 (2013) 18892–7.

[27] T. Harada, J. Swift, J. Irianto, J.W. Shin, K.R. Spinler, A. Athirasala, R. Diegmiller, P.C. Dingal, I.L. Ivanovska, and D.E. Discher, Nuclear lamin stiffness is a barrier to 3D migration, but softness can limit survival. J. Cell Biol. 204 (2014) 669–82.

[28] A.D. Stephens, E.J. Banigan, S.A. Adam, R.D. Goldman, and J.F. Marko, Chromatin and lamin A determine two different mechanical response regimes of the cell nucleus. Mol. Biol. Cell 28 (2017) 1984–1996.

[29] O. Wintner, N. Hirsch-Attas, M. Schlossberg, F. Brofman, R. Friedman, M. Kupervaser, D. Kitsberg, and A. Buxboim, A Unified Linear Viscoelastic Model of the Cell Nucleus Defines the Mechanical Contributions of Lamins and Chromatin. Adv. Sci. (Weinh) 7 (2020) 1901222.

[30] N. Zuela-Sopilniak, D. Bar-Sela, C. Charar, O. Wintner, Y. Gruenbaum, and A. Buxboim, Measuring nucleus mechanics within a living multicellular organism: Physical decoupling and attenuated recovery rate are physiological protective mechanisms of the cell nucleus under high mechanical load. Mol. Biol. Cell 31 (2020) 1943–1950.

[31] J.D. Pajerowski, K.N. Dahl, F.L. Zhong, P.J. Sammak, and D.E. Discher, Physical plasticity of the nucleus in stem cell differentiation. Proc Natl Acad Sci U S A 104 (2007) 15619–24.

[32] S. Neelam, T.J. Chancellor, Y. Li, J.A. Nickerson, K.J. Roux, R.B. Dickinson, and T.P. Lele, Direct force probe reveals the mechanics of nuclear homeostasis in the mammalian cell. Proc Natl Acad Sci U S A 112 (2015) 5720–5.

[33] Q. Zhang, A.C. Tamashunas, A. Agrawal, M. Torbati, A. Katiyar, R.B. Dickinson, J. Lammerding, and T.P. Lele, Local, transient tensile stress on the nuclear membrane causes membrane rupture. Mol Biol Cell 30 (2019) 899–906.

[34] R.B. Dickinson, A. Katiyar, C.R. Dubell, and T.P. Lele, Viscous shaping of the compliant cell nucleus. APL Bioeng 6 (2022) 010901.

[35] A. Katiyar, V.J. Tocco, Y. Li, V. Aggarwal, A.C. Tamashunas, R.B. Dickinson, and T.P. Lele, Nuclear size changes caused by local motion of cell boundaries unfold the nuclear lamina and dilate chromatin and intranuclear bodies. Soft Matter 15 (2019) 9310–9317.

[36] S. Neelam, P.R. Hayes, Q. Zhang, R.B. Dickinson, and T.P. Lele, Vertical uniformity of cells and nuclei in epithelial monolayers. Sci Rep 6 (2016) 19689.

[37] B.D. Cosgrove, C. Loebel, T.P. Driscoll, T.K. Tsinman, E.N. Dai, S.J. Heo, N.A. Dyment, J.A. Burdick, and R.L. Mauck, Nuclear envelope wrinkling predicts mesenchymal progenitor cell mechano-response in 2D and 3D microenvironments. Biomaterials 270 (2021) 120662.

[38] V.J. Tocco, Y. Li, K.G. Christopher, J.H. Matthews, V. Aggarwal, L. Paschall, H. Luesch, J.D. Licht, R.B. Dickinson, and T.P. Lele, The nucleus is irreversibly shaped by motion of cell boundaries in cancer and non-cancer cells. J Cell Physiol 233 (2018) 1446–1454.

[39] A. Jana, A. Tran, A. Gill, A. Kiepas, R.K. Kapania, K. Konstantopoulos, and A.S. Nain, Sculpting Rupture-Free Nuclear Shapes in Fibrous Environments. Adv Sci (Weinh) 9 (2022) e2203011.

[40] H.R. Thiam, P. Vargas, N. Carpi, C.L. Crespo, M. Raab, E. Terriac, M.C. King, J. Jacobelli, A.S. Alberts, T. Stradal, A.M. Lennon-Dumenil, and M. Piel, Perinuclear Arp2/3-driven actin polymerization enables nuclear deformation to facilitate cell migration through complex environments. Nat Commun 7 (2016) 10997.

[41] J. Plateau, Statique expérimentale et théorique des liquides soumis aux seulesforces moléculaires.. Gauthier-Villars. (1873).

[42] A. Janshoff, Viscoelastic properties of epithelial cells. Biochem Soc Trans 49 (2021) 2687–2695.

[43] A. Katiyar, J. Zhang, J.D. Antani, Y. Yu, K.L. Scott, P.P. Lele, C.A. Reinhart-King, N.J. Sniadecki, K.J. Roux, R.B. Dickinson, and T.P. Lele, The Nucleus Bypasses Obstacles by Deforming Like a Drop with Surface Tension Mediated by Lamin A/C. Adv Sci (Weinh) 9 (2022) e2201248.

[44] A. Katiyar, J. Zhang, J.D. Antani, Y. Yu, K.L. Scott, P.P. Lele, C.A. Reinhart-King, N.J. Sniadecki, K.J. Roux, R.B. Dickinson, and T.P. Lele, The nucleus bypasses obstacles by deforming like a drop with surface tension mediated by lamin A/C. Advanced Science In press (2022).

[45] F. Erdel, M. Baum, and K. Rippe, The viscoelastic properties of chromatin and the nucleoplasm revealed by scale-dependent protein mobility. J Phys Condens Matter 27 (2015) 064115.

[46] V.I.P. Keizer, S. Grosse-Holz, M. Woringer, L. Zambon, K. Aizel, M. Bongaerts, F. Delille, L. Kolar-Znika, V.F. Scolari, S. Hoffmann, E.J. Banigan, L.A. Mirny, M. Dahan, D. Fachinetti, and A. Coulon, Live-cell micromanipulation of a genomic locus reveals interphase chromatin mechanics. Science 377 (2022) 489–495.

[47] K.N. Dahl, S.M. Kahn, K.L. Wilson, and D.E. Discher, The nuclear envelope lamina network has elasticity and a compressibility limit suggestive of a molecular shock absorber. J Cell Sci 117 (2004) 4779–86.

[48] N. Caille, O. Thoumine, Y. Tardy, and J.J. Meister, Contribution of the nucleus to the mechanical properties of endothelial cells. J Biomech 35 (2002) 177–87.

[49] W. Helfrich, Elastic properties of lipid bilayers: theory and possible experiments. Z Naturforsch C 28 (1973) 693–703.

[50] A. Vaziri, and M.R. Mofrad, Mechanics and deformation of the nucleus in micropipette aspiration experiment. J Biomech 40 (2007) 2053–62.

[51] A. Vaziri, H. Lee, and M. Mofrad, Deformation of the cell nucleus under indentation: Mechanics and mechanisms. Journal of Materials Research 21 (2006) 2126–2135.

[52] A. Agrawal, and T.P. Lele, Geometry of the nuclear envelope determines its flexural stiffness. Mol Biol Cell 31 (2020) 1815–1821.

[53] E.M. Hatch, and M.W. Hetzer, Nuclear envelope rupture is induced by actin-based nucleus confinement. J Cell Biol 215 (2016) 27–36.

[54] H. Pan, Y.-K. Choi, Y. Liu, W. Hu, Q. Du, K. Polthier, C. Zhang, and W. Wang, Constant Mean Curvature Surfaces. ACM Transactions on Graphics 31 (2012) 85.

[55] P.-O. Persson, and G. Strang, A simple mesh generator in MATLAB. SIAM Review 46(2) (2004) 329–345.

